# Brain volumetric changes in the general population following the COVID-19 outbreak and lockdown

**DOI:** 10.1101/2020.09.08.285007

**Authors:** Tom Salomon, Adi Cohen, Daniel Barazany, Gal Ben-Zvi, Rotem Botvinik-Nezer, Rani Gera, Shiran Oren, Dana Roll, Gal Rozic, Anastasia Saliy, Niv Tik, Galia Tsarfati, Ido Tavor, Tom Schonberg, Yaniv Assaf

**Affiliations:** School of Neurobiology, Biochemistry and Biophysics, Faculty of Life Science, Tel Aviv University, Tel Aviv, Israel; Sagol School of Neuroscience, Tel Aviv University, Tel Aviv, Israel; The Strauss Center for Computational Neuroimaging, Tel Aviv University, Tel Aviv, Israel; Department of Psychological and Brain Sciences, Dartmouth College, Hanover, New Hampshire, USA; School of Psychological Sciences, Tel Aviv University, Tel Aviv, Israel; Department of Anatomy and Anthropology, Faculty of Medicine, Tel Aviv University, Tel Aviv, Israel; Division of Diagnostic Imaging, Sheba Medical Center, Tel-Hashomer, affiliated to the Faculty of Medicine, Tel Aviv University, Tel Aviv, Israel

**Author notes:** equal contribution (Ido Tavor, Yaniv Assaf).

## Abstract

The COVID-19 outbreak introduced unprecedented health-risks, as well as pressure on the financial, social, and psychological well-being due to the response to the outbreak. Here, we examined the manifestations of the COVID-19 outbreak on the brain structure in the healthy population, following the initial phase of the pandemic in Israel. We pre-registered our hypothesis that the intense experience of the outbreak potentially induced stress-related brain modifications. Volumetric changes in *n* = 50 participants who were scanned before and after the COVID-19 outbreak and lockdown, were compared with *n* = 50 control participants who were scanned twice prior to the pandemic. The pandemic provided a rare opportunity to examine brain plasticity in a natural experiment. We found volumetric increases in bilateral amygdalae, putamen, and the anterior temporal cortices. Changes in the amygdalae diminished as time elapsed from lockdown relief, suggesting that the intense experience associated with the pandemic outbreak induced transient volumetric changes in brain regions commonly associated with stress and anxiety.

## Main

Since early 2020, the world has been coping with the outbreak of the coronavirus disease 2019 (COVID-19) pandemic that infected millions with devastating numbers of deaths globally. As an initial response to the first wave of the outbreak, countries closed their borders and implemented a series of ad-hoc laws and orders to restrict the spread of the disease. Countries with major outbreaks such as China, Italy, and Spain enforced stringent restriction of movement for a limited period, referred to here as ‘lockdown’. Although lockdowns contributed to restricting the health risks of the outbreak^1^, they also had a negative impact on the social, financial and mental well-being of the general population, leading to one of the sharpest declines in economic growth over the past decades^2,3^. COVID-19 outbreak also led to high rates of stress and anxiety that were largely attributed to implications beyond actual health risk, such as the difficulties of social isolation and financial consequences of responding to the health crisis^4^. It is now evident that the indirect consequences of the pandemic affected a much larger proportion of the population, having an impact of no lesser gravity than the actual health risks that were meant to be prevented^5,6^.

In Israel, a strict lockdown period was issued from mid-March until the end of April. During its peak, most unessential businesses were closed and civilians’ movement for non-essential destinations was restricted to a radius of 100 meters from their homes. Prior to COVID-19, the country had experienced a period of peak economic prosperity^7^, which was interrupted by the outbreak, leading to unprecedented unemployment rates (reaching nearly 30% of the work-force in April 2020) and the collapse of several sectors such as aviation, tourism, and culture^8,9^. The outbreak period was characterized with acute uncertainty and increase in anxiety, regarding both the health and socioeconomic effects of the pandemic^10^.

Over the past years, several studies demonstrated the potential to detect brain plasticity using T1-weighted magnetic resonance imaging (MRI)^11–13^. The current work was initiated as a reaction to the outbreak of COVID-19 in Israel, aiming to study the structural brain plasticity in the general population following a real-life event of global scale. For this purpose, we examined *n* = 50 test group participants that were scanned with T1-weighted MRI prior to the outbreak and returned for a follow-up scan after the lockdown period (see methods for a detailed timeline of post COVID-19 follow-up scans). The structural changes of the study group (before versus after the outbreak) were compared to those of *n* = 50 control participants who were scanned twice before the COVID-19 outbreak. The unique circumstances imposed due to the COVID-19 lockdown created rare settings for a natural experiment to examine the effect of a real-world intense event on brain plasticity.

All participants were healthy, without a history of neurological or psychiatric disorders, did not show COVID-19 symptoms, and were not diagnosed carrying the virus (see the methods section for further demographic information). The hypotheses and general design of the current study were pre-registered prior to the completion of data collection and were based on a small independent pilot sample with *N* = 16 participants (*n* = 8 participants in each group). The data and analysis codes are openly shared online (project page: https://osf.io/wu37z/; preregistration: https://osf.io/k6xhn/).

## Results

Prior to their follow-up MRI scan session, we asked participants of the post-lockdown test group to fill in a short questionnaire regarding the lockdown period (see methods). Of the participants who agreed to reply, 79.6% reported they did not leave their home for non-essential needs, 79.6% met no more than 3 people, 44.9% did not meet with their parents at all, 38.8% indicated an increased feeling of anxiety following the lockdown, 34.7% anticipated that their future behavior will change after the lockdown, 46.8% reported they were concerned about their personal future well-being, 42.9% indicated that their employment status was reduced to part-time or unemployment/furlough. In an exploratory factor analysis (EFA; using Varimax rotation, see methods), we identified two main factors, explaining together 54.0% of the variance. The first factor was highly loaded with questionnaire items that described increased social isolation, and the second was mainly related to increased feelings of anxiety (Figure 1).

**Figure 1.**
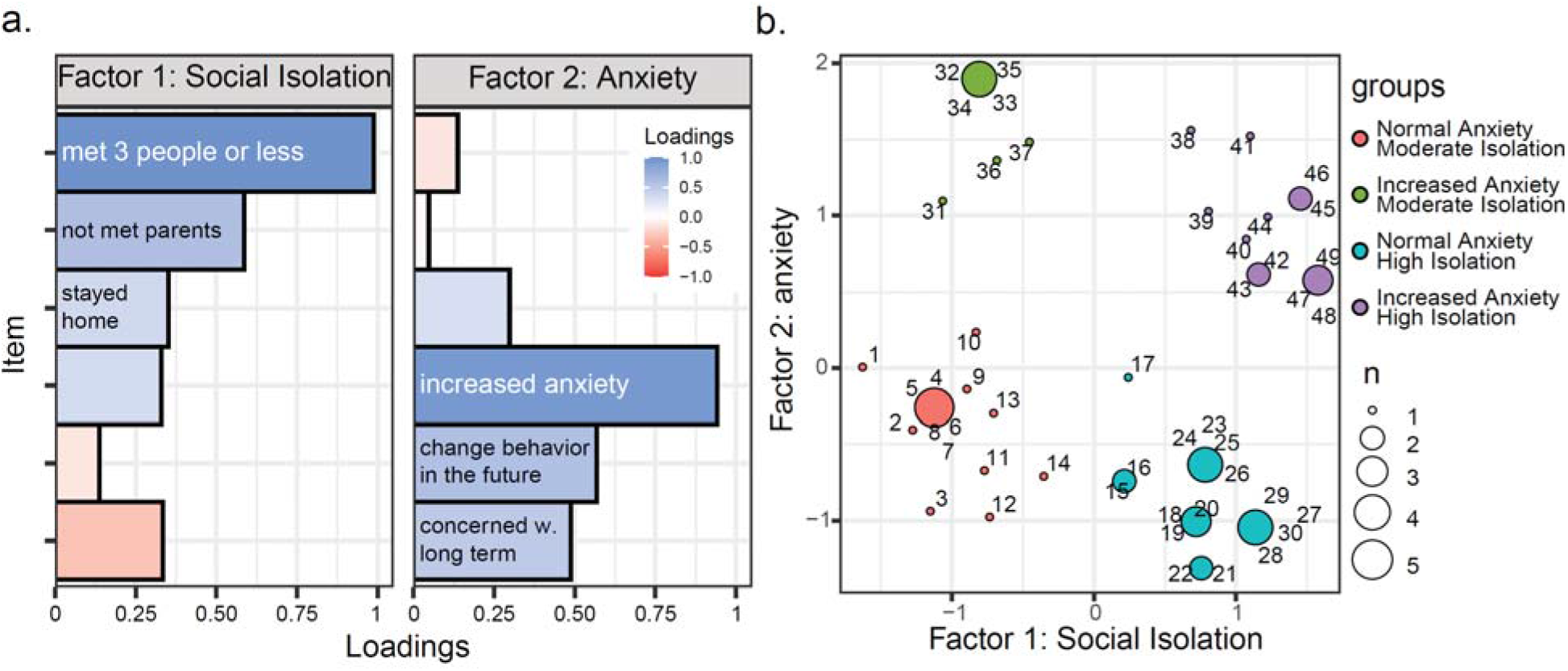
Exploratory factor analysis (EFA) of COVID-19 questionnaire. Exploratory factor analysis of the responses to the questionnaire revealed two main themes characterized the participants. **a.** The first factor (‘Social isolation’) strongly related to the item indicating meeting no more than 3 people, as well as to other two related items of avoiding meeting parents and staying at home during lockdown. An increased feeling of anxiety dominated the second factor, along with changing future behavior and concerns regarding the long-term effects. **b.** Visualization of participants dispersion across the two factors and their categorization into binary anxiety and isolation groups. Points represent unique scores. Scores’ frequency is represented by point size and participants’ indices around their corresponding data points.

Based on our pilot study results and previous studies of stress-related morphological brain changes^14–17^, we hypothesized that the focus of volumetric changes will be observed mainly in the amygdalae. T1-weighted anatomical MRI scans were used as input for deformation and surface-based morphometry (SBM) analysis using the CAT12 toolbox (http://www.neuro.uni-jena.de/cat/, University of Jena) for SPM12 (http://www.fil.ion.ucl.ac.uk/spm/software/spm12/, Wellcome Trust Centre for Neuroimaging). The brain was segmented to 58 regions based on the cortical and subcortical nuclei classifications of the Hammers atlas (Hammers et al., 2003). Following surface reconstruction, each participant’s individual gray matter volume was estimated for each of the 58 anatomically defined regions of interest (ROIs). This procedure accounted for the longitudinal nature of the data, performing the analysis on both scans simultaneously. To avoid voxel-based multiple comparisons, we performed a region-based analysis (following surface projection to the Hammer atlas) and corrected for multiple comparisons using the Benjamini-Hochberg correction^18^ to control for false discovery rate (FDR; using *p* < 0.05 following correction threshold). Validation of the pipeline was performed using simulated data and by comparing the results with other software (see methods).

Using a linear mixed model, we examined volumetric changes, testing for regions with stronger changes for the test group compared to the control group. Examining the interaction effect of session (baseline versus follow-up scans) and experimental group (test versus control) revealed ten anatomical brain regions (composed of bilateral five unique regions in both hemispheres) in which volumetric increases were observed uniquely for the test group (Table 1 and Figure 2). Most prominently, as we expected and pre-registered, we found a robust effect in the bilateral amygdalae. We also observed a significant effect bilaterally in the putamen, and in three anatomical regions within the ventral anterior temporal cortex adjacent to each other, namely in the medial part of the anterior temporal lobe, the fusiform gyrus, and the parahippocampal gyrus. To examine the spatial distribution within significant ROIs, we performed an additional post-hoc voxel-based analysis, which allowed us to visualize the changes within the significant ROIs (Figure 2a). Examining the post-hoc voxel-based results revealed that volumetric changes occurred throughout the entire surface of bilateral amygdalae, while in the putamen the effects occurred mainly in the dorsal area. In the ventral anterior temporal cortices, large connected clusters of volumetric change spanned throughout the three adjacent temporal ROIs, thus suggesting that the three ROIs shared a similar origin. To ensure that the reported effects originated from volumetric changes in the test group following the COVID-19 outbreak and its related lockdown period, we tested for ROIs where the significant interaction effect was accompanied by a significant effect for the test group but not for the control group, and was consistent beyond baseline scans effect or measurement protocol (see methods).

**Table 1.** Surface based morphology analysis results

| Region | Hemi-sphere | Interaction estimate<br>(95% CI) | Interaction *p*<br>(FDR adj.) | Test group session estimate<br>(95% CI)<sup>a</sup> | Test group session *p*<br>(FDR adj.) | Change from baseline *M* (*SE*)<sup>b</sup> |  |
| --- | --- | --- | --- | --- | --- | --- | --- |
|  |  |  |  |  |  | Test | Control |
| Amygdala | Left | 0.09<br>[0.05, 0.13] | 2.4E<sup>-5</sup><br>(0.001) | 0.08<br>[0.05, 0.11] | 9.8E<sup>-6</sup><br>(2.1E<sup>-4</sup>) | 4.92%<br>(1.06%) | -0.56%<br>(0.6%) |
|  | Right | 0.08<br>[0.03, 0.13] | 0.003<br>(0.030) | 0.08<br>[0.05, 0.11] | 1.6E<sup>-5</sup><br>(2.3E<sup>-4</sup>) | 4.47%<br>(1.07%) | 0.07%<br>(0.86%) |
| Putamen | Left | 0.19<br>[0.09, 0.29] | 4.1E<sup>-4</sup><br>(0.006) | 0.13<br>[0.06, 0.2] | 4.0E<sup>-4</sup><br>(0.002) | 3.31%<br>(0.83%) | -0.5%<br>(0.7%) |
|  | Right | 0.17<br>[0.08, 0.26] | 2.4E<sup>-4</sup><br>(0.005) | 0.14<br>[0.08, 0.2] | 1.1E<sup>-5</sup><br>(2.1E<sup>-4</sup>) | 3.31%<br>(0.67%) | -0.01%<br>(0.68%) |
| Anterior temporal lobe (medial part) | Left | 0.25<br>[0.12, 0.38] | 1.8E<sup>-4</sup><br>(0.005) | 0.15<br>[0.07, 0.23] | 4.7E<sup>-4</sup><br>(0.003) | 2.82%<br>(0.75%) | -0.7%<br>(0.76%) |
|  | Right | 0.21<br>[0.07, 0.35] | 0.004<br>(0.030) | 0.15<br>[0.05, 0.25] | 0.004<br>(0.023) | 2.93%<br>(1.07%) | -0.37%<br>(0.7%) |
| Parahippocampal gyrus | Left | 0.09<br>[0.03, 0.15] | 0.006<br>(0.035) | 0.04<br>[0, 0.08] | 0.029<br>(0.085) | 1.22%<br>(0.55%) | -0.5%<br>(0.57%) |
|  | Right | 0.11<br>[0.04, 0.18] | 0.003<br>(0.030) | 0.08<br>[0.03, 0.13] | 0.002<br>(0.009) | 2.05%<br>(0.61%) | -0.08%<br>(0.54%) |
| Fusiform gyrus | Left | 0.08<br>[0.03, 0.13] | 0.007<br>(0.036) | 0.06<br>[0.03, 0.09] | 3.8E<sup>-4</sup><br>(0.003) | 1.78%<br>(0.53%) | -0.4%<br>(0.56%) |
|  | Right | 0.11<br>[0.04, 0.18] | 0.002<br>(0.022) | 0.05<br>[0, 0.1] | 0.044<br>(0.111) | 1.36%<br>(0.64%) | -0.72%<br>(0.57%) |

<sup>a</sup> Session estimate examined the effect of baseline versus follow-up scan in the post-lockdown test group. This parameter was used to validate that the interaction effect observed between the group stemmed from a robust effect in the test group (see methods).
<sup>b</sup> Volumetric change normalized to baseline scan (difference/ baseline).

**Figure 2.**
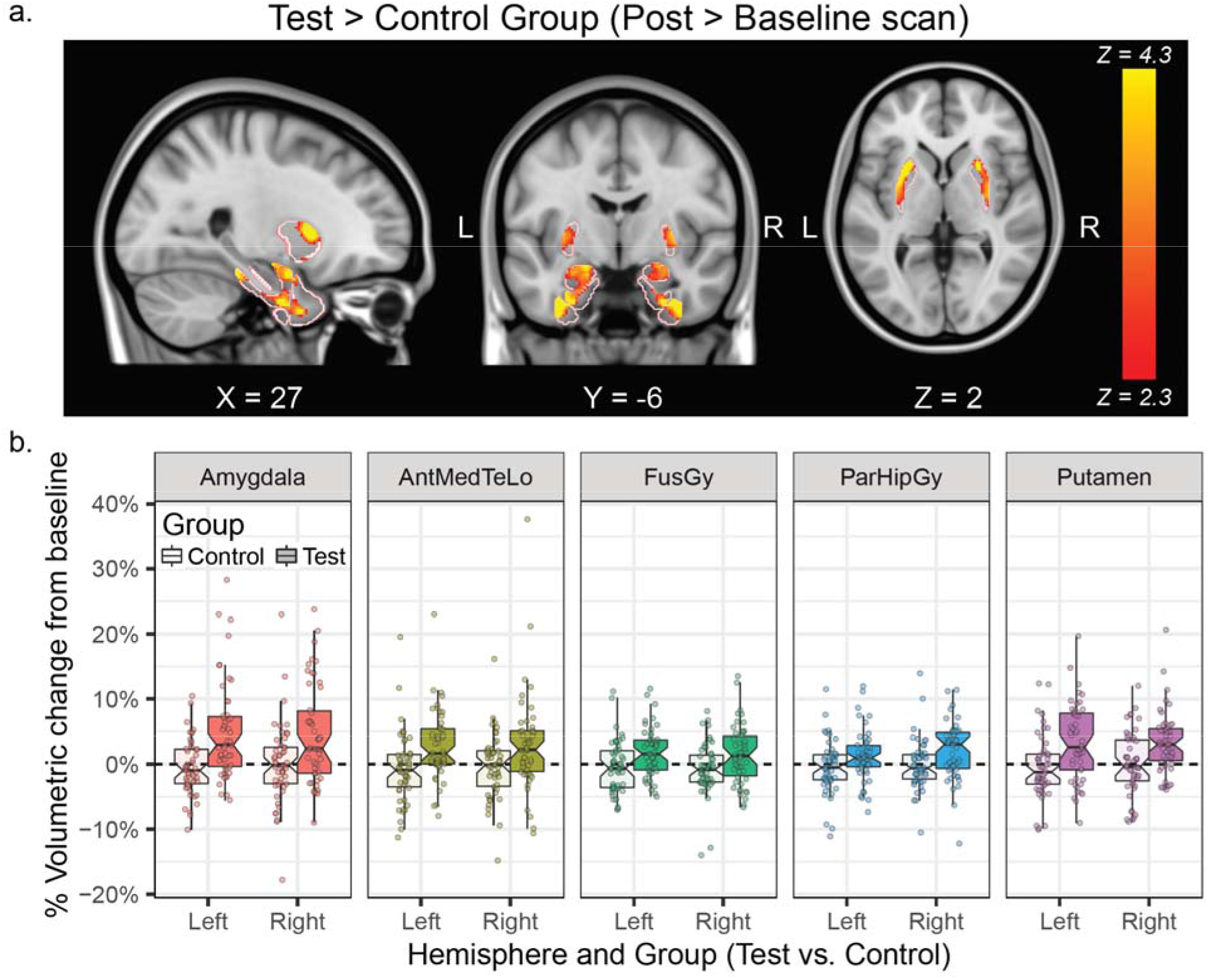
Volumetric changes results. An interaction effect for time (baseline versus follow-up scan) and group (test versus control) was evaluated on segmented surfaces in an SBM analysis. Significant interaction effects were observed bilaterally in the amygdala and putamen ROIs, as well as in three ventral temporal cortical ROIs. **a.** To examine spatial patterns within the identified ROIs, a post-hoc voxel-based analysis was conducted within each ROI mask. Light red contours represent segmentation borders of the ROIs. **b.** Individual distribution of the results in the control group (light colors) and test group (dark colors). For better visualization, units were normalized to baseline (difference/baseline) and presented in percentage units. Box-plot center, hinges, and whiskers represent the median, quartiles, and from the hinges, respectively. A notch of represent an estimated 95% confidence interval for comparing medians. Dots represent individual participants. Abbreviated ROI names: AntMedTeLo = anterior temporal lobe (medial part); FusGy = fusiform gyrus, ParHipGy = Parahippocampal gyrus.

To evaluate and control for the effect of time between scans and time from lockdown, we included in the model two additional covariates - the time between scans (TBS; which was generally longer for the test group) and time following lockdown (TFL; see methods for more details). The two covariates were not correlated with each other in our test group sample (*r* = −0.106, *t*(48) = −0.74, *p* = 0.463). Our reported regions demonstrated significant volumetric change above and beyond these covariates. After FDR correction, no region showed an effect of TBS. However, we did find a negative effect of TFL in the two amygdalae ROIs and the left fusiform gyrus, suggesting that the volumetric changes in these regions moderated as time following lockdown elapsed. Based on these results, we estimated the time to decay as the estimated number of days from lockdown until volumetric changes returned to normal levels, similar to those of the control group (left amygdala: *β*_TFL_ = −0.41, *t*(47) = −3.1, *p* = 0.003, *p*_adj_. = 0.048, time to decay = 95 days; right amygdala: *β*_TFL_ = −0.54, *t*(47) = −4.38, *p* = 6.7E^−5^, *p*_adj_. **=** 0.002, time to decay = 83 days; left fusiform gyrus: *β*_TFL_ = −0.54, *t*(47) = −4.44, *p* = 5.5E^−5^, *p*_adj_. = 0.002, time to decay = 82 days; Figure 3).

**Figure 3.**
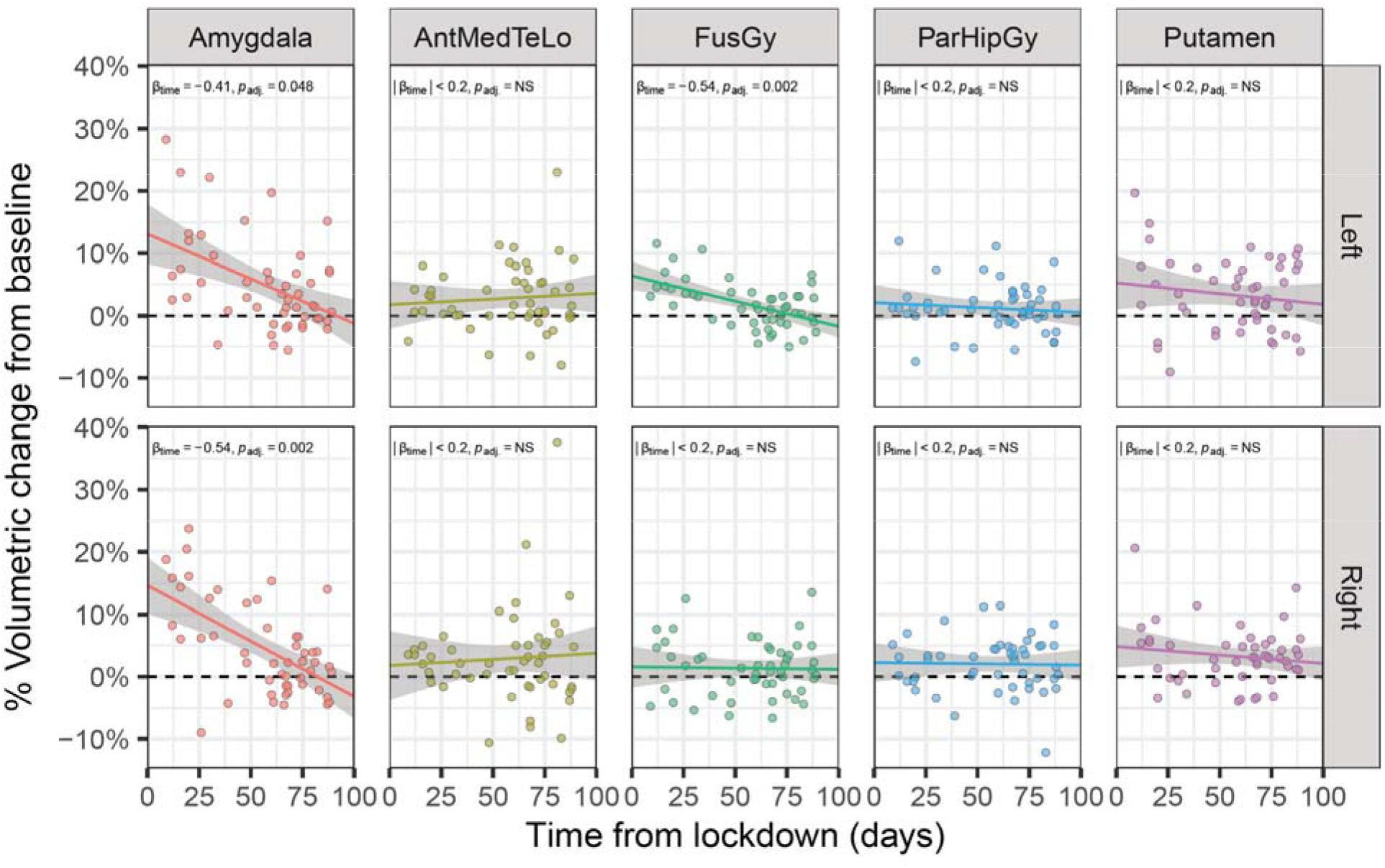
Time following lockdown effect on volumetric changes. The time from lockdown relief until the follow-up scan session (TFL) was added as an addition covariate to the model, revealing significant effect in the two amygdalae and left fusiform gyrus. Points represent individual participants in the post-lockdown test, *p*-values were FDR adjusted for multiple comparisons. Abbreviated ROIs: AntMedTeLo = anterior temporal lobe (medial part); FusGy = fusiform gyrus, ParHipGy = Parahippocampal gyrus.

In addition to the volumetric change effect reported here, we also performed an exploratory analysis aimed to detect pattern of correlation change between all tested ROIs, and how do these changes differ between the two groups (see supplementary results). We also performed additional exploratory analyses to examine the association of volumetric changes and the reported experience during lockdown. We found no strong association between the two, as reported in the supplementary results.

## Discussion

Our study demonstrates that volumetric change patterns in the brain occurred following the COVID-19 initial outbreak period and restrictions in a sample of healthy participants, who were not somatically affected by the pandemic. While previous studies demonstrated brain plasticity using T1-weighted MRI following planned interventions^11–13^, the current work outstands in its unique demonstration of stark structural brain plasticity following a major real-life event.

Our findings show neural changes that were not caused directly due to COVID-19 infection, but rather related to the societal effect. We show volumetric increase in gray matter in the amygdalae, putamen and ventral anterior temporal cortices. The changes in the amygdalae showed a temporal-dependent effect, related to the time elapsed from lockdown but not the duration from the baseline scan. It should be noted that although lockdown restrictions had initially reduced infection rates in Israel, just one month after the lockdown was lifted, the number of infected cases started to rise again and reached higher number of active infected cases by the end of data collection, compared with the peak numbers during the actual lockdown period (approximately 2,000 daily new cases by the end of July versus under 750 new daily cases during the peak of the lockdown period in April^19^, see methods and Figure 4). This suggests that the effects observed in the current study are less likely to be attributed to the concrete health risks of COVID-19, but rather to the first wave of the outbreak, characterized with perceived uncertainty and substantial unexpected changes in everyday life.

**Figure 4.**
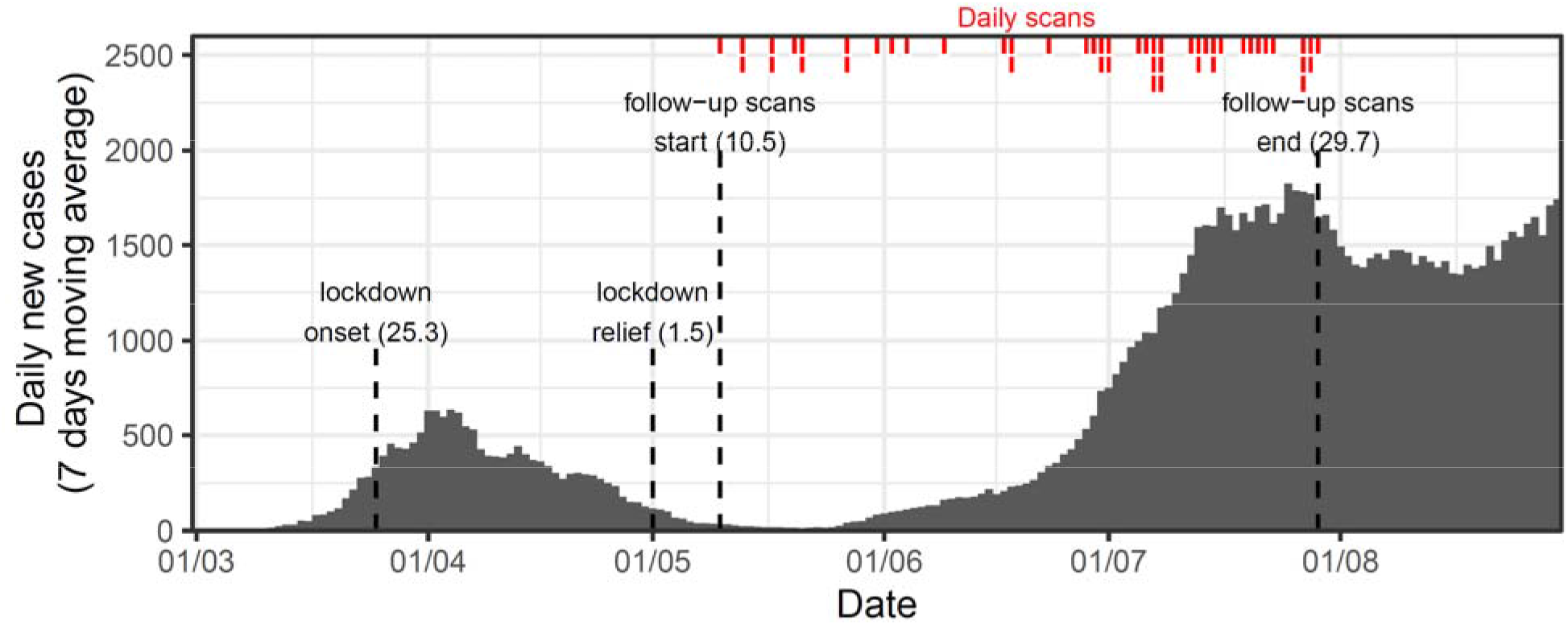
Study timeline and outbreak data On February 21^st^, 2020, the first COVID-19 case in Israel was recorded. Daily new cases we**re** smoothed using 7-days moving average. Data were retrieved and modified based on the Isra**eli** Ministry of Health reports^19,23^. A lockdown was issued on March 25^th^, which was gradually released until the removal of the 100-meter restriction on May 1^st^, marking lockdown onset and relief, respectively (shorter vertical dashed line). MRI data of the test group were collected between May 10^th^ to July 29^th^ (longer vertical dashed line). Red bars on top represent the number of participants scanned for the study each day.

The current study was in many aspects unplanned; therefore, we are left with only partial answers as to which specific components of the COVID-19 outbreak led to the neural changes observed in the healthy participants that took part in our study. The involvement of the amygdala may suggest that stress and anxiety could be the source of the observed phenomenon, due to its well-recorded functional and structural associations ^14–17,20–22^. Nevertheless, it is hard to draw clear conclusions as many aspects of life have changed in this time period, and could have potentially affected different regions in the brain – from limiting social interactions, increased financial stress, changes in physical activity, work routine, and many more. The limited behavioral data collected in the current study did not provide a strong connection to the imaging results, and thus future work could try to better address the complex brain-behavioral associations in this real-life experience. Nonetheless, our findings show that healthy young adults, with no records of mental health issues, were deeply affected by the outbreak of COVID-19. We suggest that policy makers take into consideration the impact of their actions on the general well-being of the population they seek to help, alongside the efficacy of disease prevention.

## Methods

### Data and codes availability

Our sample size, hypotheses and analyses plan were pre-registered on the Open Science Framework (OSF), soon after data collection began, but prior to completion of the data collection and data analysis (project page: https://osf.io/wu37z/; preregistration: https://osf.io/k6xhn). All behavioral data, processed imaging data, and analysis codes are shared on the OSF project page. Uncorrected and small-volume corrected statistical maps of the voxel-based results described in the current work are available at https://neurovault.org/collections/8591/.

### Participants

The study included two groups: A test group scanned before and after COVID-19 lockdown, and a control group, scanned twice before COVID-19 lockdown. All participants had no background of neurological disorders, did not show symptoms for COVID-19 and were not diagnosed as carriers of the virus. The study was approved by the ethics committee of Tel Aviv University and institutional review board (IRB) at the Sheba Tel-Hashomer medical center. Since the IRB protocol allowed us to scan the participants several times over a long period of time, we were able to collect the data from participants who were scanned prior to COVID-19 outbreak and invite them back for a follow-up scan as part of the longitudinal study they have agreed to take part in. Participants received monetary compensation for their time and gave their informed consent to take part in a longitudinal experiment aimed to examine brain plasticity across several sessions, which was initially not directly related to COVID-19 outbreak.

The test group comprised of *n* = 50 participants who were scanned before and after COVID-19 lockdown (Δ Time between scans: *M* = 309.3, *SD* = 207.5, range = 67 - 1460 days; Age: *M* = 30.1, *SD* = 6.65, range = 21 - 48; Females: n = 20, prop. = 40%). The lockdown period began on March 25^th^ and was gradually relieved throughout late April. We mark here May 1^st^ as the lockdown relief date, as on this day an issued 100-meters movement limit for non-essential needs was lifted. The test group data collection started as soon as lockdown relief took place, for a period of approximately 3 months, until the end of July, 2020 (Δ Time from lockdown relief: *M* = 57 days, *SD* = 24.62, range = 9 - 89 days; see Figure 4 for the study timeline).

As a control group, we used the data of *n* = 50 participants who were scanned twice using a similar protocol before COVID-19 lockdown (Δ Time between scans: *M* = 126.7, *SD* = 190.4, range = 21 - 886 days; Age: M = 27.3, *SD* = 5.63, range = 19 - 42; Females: n = 23, prop. = 46%).

The minimal sample size was determined and pre-registered (https://osf.io/uktsn), based on a 80% power analysis conducted using R ‘pwr’ package^24^, on a pilot study with *N* = 16 participants (*n* = 8 in each group). We decided to collect a minimum of *n* = 37 participants which should provide 80% to detect the group and session interaction effect with α = .05, within both the left and right amygdala, under the assumption that it would be difficult to complete the sample due to COVID-19 limitations. Eventually, thanks to further relief in COVID-19 restrictions, we were able to extend the sample size to *n* = 50 in each group.

The results remain generally consistent, even when the data included only the first *n* = 37 participant; demonstrating significant effects in the bilateral amygdalae, putamen, parahippocampal gyrus and the left anterior temporal lobe. In this smaller sample, we did not find significant effects in the right anterior temporal lobe, nor in the fusiform gyrus. We also found volumetric increase effects that were not identified using the full sample in the left nucleus accumbens, left cuneus, and left insula.

### Imaging data

#### Acquisition protocol

Imaging data were acquired using a 3T Siemens Prisma scanner, with a 64-channel head coil. For the structural data, T1w high resolution (1-mm^3^) whole brain images were acquired with a magnetization prepared rapid gradient echo (MPRAGE) pulse sequence with repetition time (TR) of 2.53s, echo time (TE) of 2.88ms, flip angle (FA) = 7°, field-of-view (FOV) = 224 × 224 × 208 mm, resolution = 1 × 1 × 1. Since for this project we combined data from different labs, some of the participants had their scans aligned to the anterior commissure - posterior commissure (AC-PC) line, while others were scanned with a 30° angle (50% of test group participants and 80% of control group participants).

Some participants were also scanned with diffusion-weighted echo-planar imaging (DW EPI) sequence and some with functional gradient-echo EPI (GE EPI) in a resting state scan. The analyses of these scans are beyond the scope of the current study.

#### Data processing and analysis

The T1-weighted MPRAGE anatomical scans were used for a surface-based morphometry (SBM) analysis. From the images we estimated the pial and inner surfaces of the cortex and projected those into the Hammers atlas (Hammers et al., 2003). Data were preprocessed in SPM12 (http://www.fil.ion.ucl.ac.uk/spm/software/spm12/, Wellcome Trust Centre for Neuroimaging) and SPM based CAT12 (Computational Anatomy Toolbox 12; http://www.neuro.uni-jena.de/cat/, University of Jena) extension. We deployed the CAT12 surface-processing pipeline, which includes skull striping, a denoising filter^25^ projection-based thickness estimation^26^, partial volume correction, and spatial normalization to MNI space. Prior to the preregistration of the protocol, we validated the reliability of our surface-based image processing pipeline, by simulating artificial activity in the amygdala, and using FreeSurfer based analysis of cortical regions (See supplementary materials for further details).

Surface-based volumetric data of cortical and subcortical regions were segmented based on the Hammers Atlas, segmenting the volumetric data into 58 anatomically defined regions. Ten additional ROIs of non-gray matter or non-cerebral structures (ventricles, white matter, brain-stem and cerebellum ROIs) were excluded from statistical analyses. To evaluate the effect of lockdown on volumetric imaging data we ran a mixed linear model on the data within each one of the 58 anatomical regions, examining the effect of session (baseline versus follow-up scan) and group interaction (test versus control), controlling for time between scans (TBS) and time following lockdown (TFL) covariates. Both covariates were mean centered before they were added to the model. To include TFL covariate in the same model with control group participants, for which TFL was irrelevant, control groups’ participants were assigned with the mean TFL; thus, their data was not used to evaluate the effect of TFL.

To identify our regions of interest we only included regions that showed both a significant interaction effect (i.e. the volumetric difference between the two sessions was significantly different for the test and control group), and a significant session effect within the test group (i.e. a significant difference between the two sessions for the test group). Results were corrected for multiple comparisons using the false discovery rate (FDR) correction^18^, based on the number of brain regions tested. Following the analysis pipeline, we identified ten significant ROIs: bilateral amygdalae, putamen, parahippocampal gyrus, the medial part of the anterior temporal lobe, and the fusiform gyrus.

An additional ROI of the right inferior and middle temporal gyri showed a significant interaction effect (interaction estimate = 0.17, 95% CI = [0.05, 0.29], *p* = 6.1E^−3^, *p*_adj_. = 0.035), however examining the test group separately, we could not identify a significant session effect (session estimate = 0.04, 95% CI = [−0.02, 0.10], *p* = 0.259, *p*_adj_. = 0.424). Therefore, it is harder to interpret that this interaction effect stemmed from the test group. Thus, we did not include this ROI as one of our significant ROIs. A less robust session effect within the test group was also observed for the left parahippocampal gyrus (session estimate = 0.04 [0.00, 0.08], *p* = 0.029, *p*_adj_. = 0.085) and right fusiform gyrus ROI (session estimate = 0.05 [0.00, 0.10], *p* = 0.044, *p*_adj_. = 0.111), however as these ROIs demonstrated strong interaction effects and their session effects were significant before FDR correction, we decided to report them together with the other significant regions. A similar procedure was used in the pre-registration (i.e. including only regions that showed a significant interaction and an effect in the test group); however, in the analysis of the pilot for the pre-registration, we used uncorrected results due to the small sample size.

To examine the spatial distribution of our effect within significant ROIs, that were identified with the SBM analysis, we performed an additional post-hoc voxel-based analysis. We projected the data on a voxel-based map, effectively examining which voxels demonstrated an interaction effect. Then, we used anatomical masks of ROIs which were found to be significant in the SBM analysis, to visualize the results within these regions, similarly to a small-volume correction analysis (Figure 2a).

### Behavioral data

#### Data collection

To evaluate participants’ experience in the peak days of the COVID-19 outbreak, we asked them to think back on their experience during this time and fill out a 7-items questionnaire regarding their experience of the COVID-19 lockdown (see Table 2 for a description of the items). The questionnaires were filled out after the initiation of the study, when the lockdown’s stringent 100-meters limitation was lifted, thus the results represents the participants’ recalled experience of the lockdown. Most participants filled out the questionnaire on the day of the post-lockdown scan session, some filled it a few days before their second scanning session. A total of *n* = 76 participants filled out the COVID-19 questionnaire and comprised the potential pool of test group participants for the current study, out of which *n* = 50 were sampled and scanned. One participant was scanned but did not complete the questionnaire, therefore this participant’s behavioral data were not used and analyses of the questionnaire were based on *n* = 49 valid participants.

**Table 2.** COVID-19 lockdown questionnaire

| Question | Possible Answers (prop.) | Binary outcome (prop.) |
| --- | --- | --- |
| 1. Did you stay home during the lockdown, except for essential needs / did not leave at all? | 0 - no (20.4%)<br>1 - yes (79.6%) | 0 - no (20.4%)<br>1 - yes (79.6%) |
| 2. Did the lockdown increase your feeling of anxiety? | 0 - no (61.2%)<br>1 - yes (38.8%) | 0 - no (61.2%)<br>1 - yes (38.8%) |
| 3. With how many people did you meet during the lockdown (including people you are living with at home)? | 0 – none (0%)<br>1 - up to three people (57.1%)<br>2- up to five people (22.4%)<br>3 - up to ten people (20.4%) | 0 - more than three (20.4%)<br>1 - up to three (79.6%) |
| 4. Do you think your behavior will change following the lockdown? | 0 - no (65.3%)<br>1 - yes (34.7%) | 0 - no (65.3%)<br>1 - yes (34.7%) |
| 5. How did your meeting with your parents' routine look like during the lockdown? | 0 - same as before the lockdown (34.7%)<br>1 - with precaution measurements: distancing, mask, etc. (20.4%)<br>2 - did not meet at all (44.9%) | 0 - as before or with precautions (55.1%)<br>1 - did not meet at all (44.9%) |
| 6. What was your employment status during the lockdown? | 0 - same as before lockdown (28.6%)<br>1 - full time working from home (28.6%)<br>2 - part time working from home (8.2%)<br>3 - Furlough / unemployed (34.7%) | 0 - unemployed / part time (42.9%)<br>1 - same as before / full time from home (57.1%) |
| 7. How concerned are you with the long-term effect of the lockdown, regarding yourself? | Ordinal scale<br>1 – not at all (28.6%)<br>2 (24.5%)<br>3 (30.6%)<br>4 (14.3%)<br>5 – very concerned (2%) | 0 - low, score 1-2 (53.1%)<br>1 - moderate-high, score 3-5 (46.9%) |

#### Exploratory factor analysis (EFA)

Responses to the lockdown questionnaire were coded into binary responses, based on the sample median, splitting the sample into relatively similar sized groups for each item (Table 2). To identify the main themes in the questionnaire, which could be correlated with the imaging data, we performed an exploratory factor analysis (EFA) on the binarized data, using the “psych” R package^27^.

Since EFA require large number of participants, we used the data from all available *n* = 75 participants who completed the questionnaire. Kaiser-Meyer-Olkin (KMO) factor adequacy test revealed that the ‘employment’ item had a very low measure of sampling adequacy (MSA; employment item’s MSA = 0.26; which is far below the suggested minimal MSA of 0.5^28^) for EFA. The ‘employment’ was also not loaded to any of the factors; therefore, we removed it from the final EFA model. Overall, even after removing low KMO item, our data was found to be weakly appropriate for factor analysis, with overall MSA = 0.48. Thus, considering the small sample size and low fit of the data to EFA, the results of the analysis should be interpreted with caution.

We performed polychoric correlations based EFA, which is suitable for binary variables, with two factors and Varimax rotation, assuming orthogonality between the factors. Our selection of number of factors was based on visual inspection of scree plot of eigenvalues, as well as by comparing actual data to simulations of random data matrices. Using Oblimin rotation, which allows for correlation between the factors, revealed very low correlation (*r* = 0.08), suggesting that Varimax rotation was an appropriate choice for our model.

To examine the association of the behavioral data with the volumetric changes, while maintaining relatively limited number of multiple comparisons, we used the two factors for these analyses instead of each of the items. These two factors’ scores for each participant were extracted and correlated with the change in gray matter volumetric data in our regions of interest (see supplementary materials for results of this exploratory analysis).

## Acknowledgements

This work was supported by the Israeli Science Foundation granted to Yaniv Assaf (ISF 1314/15), Tom Schonberg (2004/15), and Ido Tavor (ISF 1603/18). Tom Salomon was supported by the Nehemia Levtzion fellowship and the Fields-Rayant Minducate Learning Innovation Research Center.

## Author contributions

T.Sa. wrote the manuscript with Y.A, assisted with the study design and analyzed the data. Data was collected by A.C., R.B.-N., R.G, S.O, G.R., D.R, and A.S. Free-surfer analysis was made by G.B-Z. N.T performed VBM validation analysis. D.B. designed the scanning protocols. G.T. provided support and medical supervision. I.T., T.Sc. and Y.A. conceived the study, wrote the manuscript and supervised the study. Y.A performed the preprocessing and analysis of imaging data. All Author contributed intellectually and reviewed the manuscript.

## Supplementary materials

### Validation using different statistical modeling

In our work, we found volumetric change effect following COVID-19 outbreak and lockdown. Our principle statistical analysis was based on the differences in volume values of the baseline and follow-up scans of each participant. To easily compare the effect in each brain region, we report and illustrate the change effect in units of percent change: 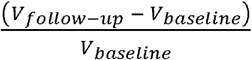.

Supplementary Figure 1, provides a visualization of the results presented in Figures 2b and 3, using the original units, without normalizing for baseline scan unites 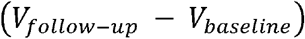.

**Supplementary Figure 1.**
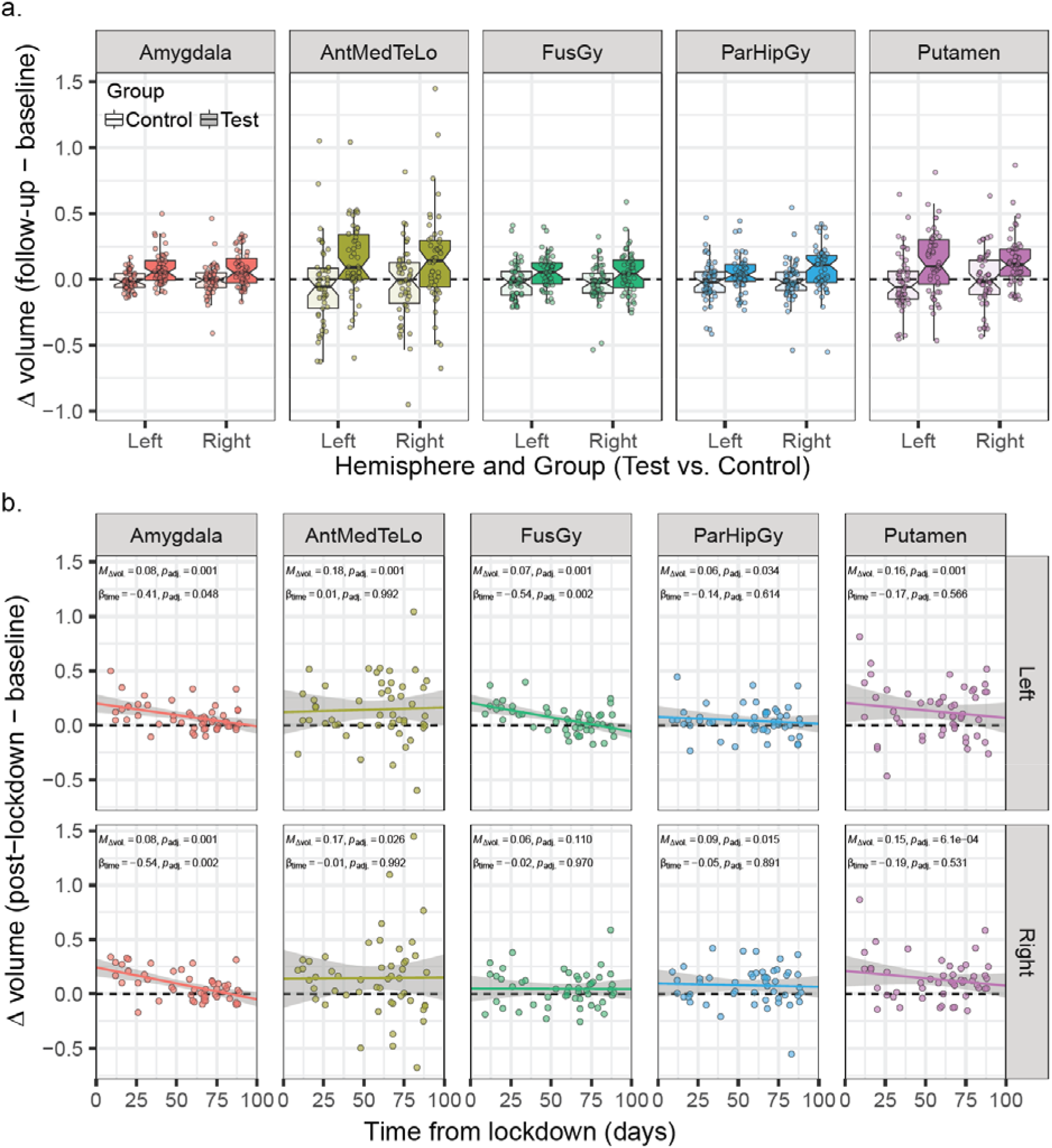
Volumetric change effects not normalized to baseline. Visualization of the reported volumetric change effects, without normalizing the data to baseline volume values. **a.** distribution of volumetric change values across the 10 signifiacant ROIs, as presented in Figure 2b. **b.** The association of volumetric change effect and time from lockdown release, as presented in Figure 3.

To further corroborate the validity of our results, we examined the volumetric change effect through several additional confirmatory perspectives, which could help exclude potential confounds. First, we examined whether the difference effect might be related to baseline scan values. To make sure that the volumetric change effect that we observed in the test group is robust above and beyond initial baseline scan, we examined a linear model with the follow-up scan volume as our dependent variable and the baseline scan as an additional covariate. The results showed consistent interaction effect. In all 10 significant ROIs, even when baseline scans explained a significant portion of the variance in follow-up scans, the difference effect between the two groups remained significant above and beyond the effect of baseline scan (Supplementary figure 2).

**Supplementary Figure 2.**
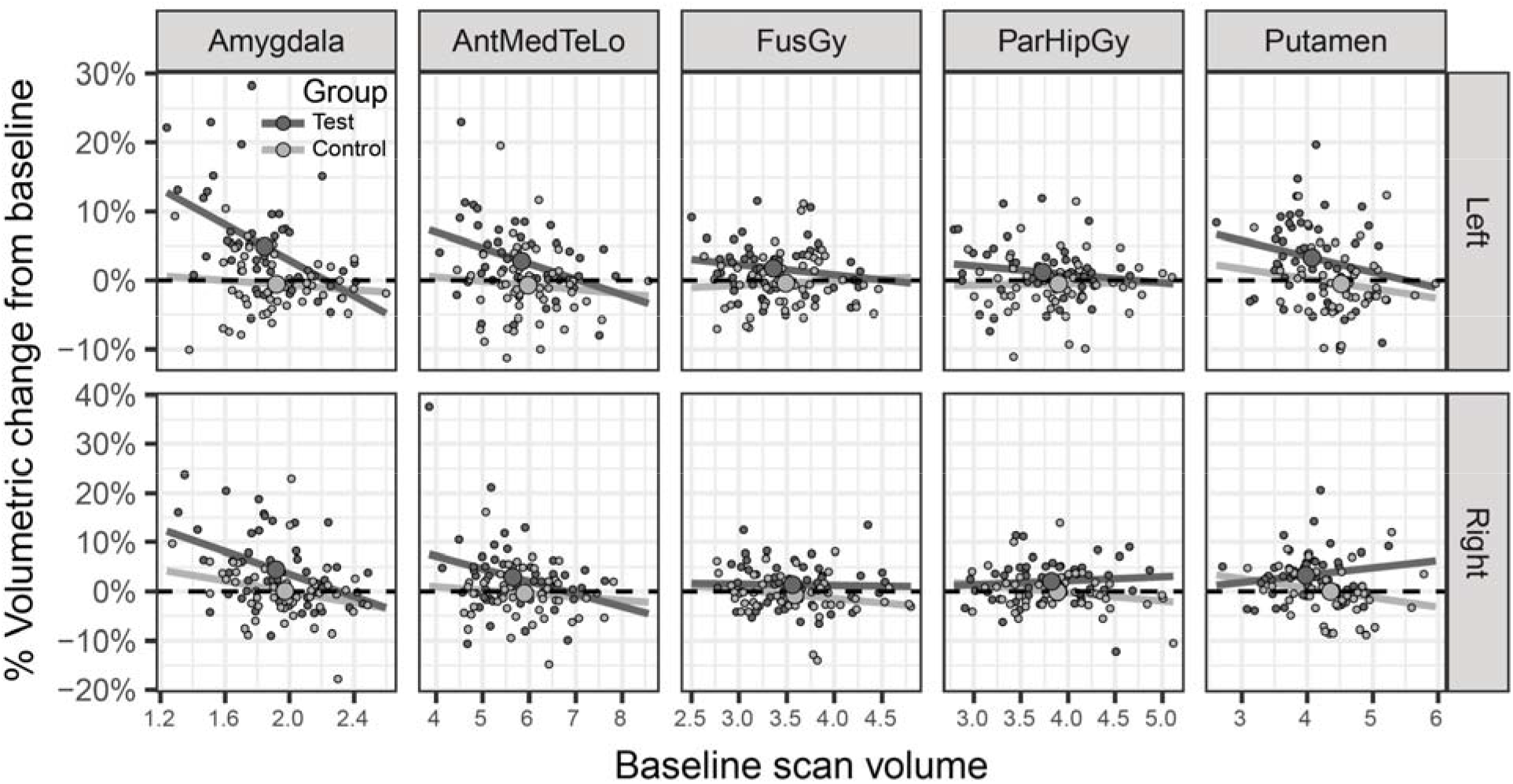
Volumetric change and baseline scan Volumetric change (difference between follow-up and baseline scans, relative to baseline) of the test group (dark grey) and control group (light grey), plotted against the volumetric values measured at baseline. Small dots represent individual participants, large dots represent the groups means. A significant interaction effect was observed above and beyond the variability explained by baseline scan values, as test group participants overall showed more robust volumetric changes than the control participants.

In another model, we also included the gender and the scan angle (AC-PC line versus 30° angle from the AC-PC) as additional independent variables. Both gender and scan angle had no significant contribution beyond other independent variables (*p*_adj_. > 0.05 in all reported ROIs). Adding gender had no effect on the reported results. However, the scan angle was correlated with the group and the TFL variables, such that a larger proportion of control-group participants were scanned with the 30°angle (80% versus 50% in the test group); and in the test group, participants who were scanned with the 30° angle were mostly scanned in the later days of data collection (TFL and scan-angle correlation: *r* = 0.615, *t*(48) = 5.41, *p* = 2.0E^−6^). The addition of the scan angle to the model therefore had an impact on the significance of other effects: (1) all reported TFL effects became non-significant after FDR-correction; and (2) while no additional regions became significant for the group effect, the FDR-corrected p-value dropped just slightly below the statistical significance threshold for the right amygdala (*p*_adj_. = 0.058), right anterior medial temporal lobe (*p*_adj_. = 0.058), right parahipocampal gyrus (*p*_adj_. = 0.058), left parahipocampal gyrus (*p*_adj_. = 0.074), right fusiform gyrus (*p*_adj_. = 0.058) and left fusiform gyrus (*p*_adj_. = 0.074). The other regions remained significant (left amygdala *p*_adj_. = 0.003, left anterior medial temporal lobe *p*_adj_. = 0.007, left putamen *p*_adj_. = 0.010, and right putamen *p*_adj_. = 0.007). These results suggest that the scan angle had a slight impact on the significance of the effects, probably due to poor balance of the condition across groups and time. Although some fail to exceed the stringent statistical threshold after FDR by a small margin, overall the results remain fairly consistent.

We further validated that our choice of measurement to analyze (difference of follow-up – baseline scans) had no significant impact on our conclusions. In another analysis, we compared the log-ratio between scans as our dependent variable, wherein log(follow-up/baseline) under the null hypothesis is expected to be 0. Using the same analysis process with log-ratio scores as the dependent variable resulted in the same 10 significant ROIs. Thus, our additional analysis suggested that our reported results are robust to potential biases due to baseline scans and choice of statistical analysis procedure.

### Imaging pipeline validation

To validate the imaging processing protocol we used two approaches, before data collection was completed (at the time of the pre-registration finalization). As many surface-based software focus on analysis of cortical surfaces, rather than subcortical regions, we aimed to validate that our SBM protocol using CAT12 could reliably identify subcortical morphological changes, such as the ones we observed in the current study within the amygdala and putamen. To test the detection capabilities of our protocol, we generated simulated data with volumetric changes in the amygdala and ran the CAT12vpipeline on the simulated data. A volumetric increase in the amygdala was simulated using a hand-drawn polygon mask, surrounding the left amygdala on the original T1w images of 10 participants. Within this polygon 3D mask, the signal intensity was artificially changed. Following this procedure, both the original and modified T1w images underwent the same CAT12 pipeline. We were able to identify the simulated volumetric changes within the subcortical amygdala nuclei (see supplementary Figure 3).

**Supplementary Figure 3.**
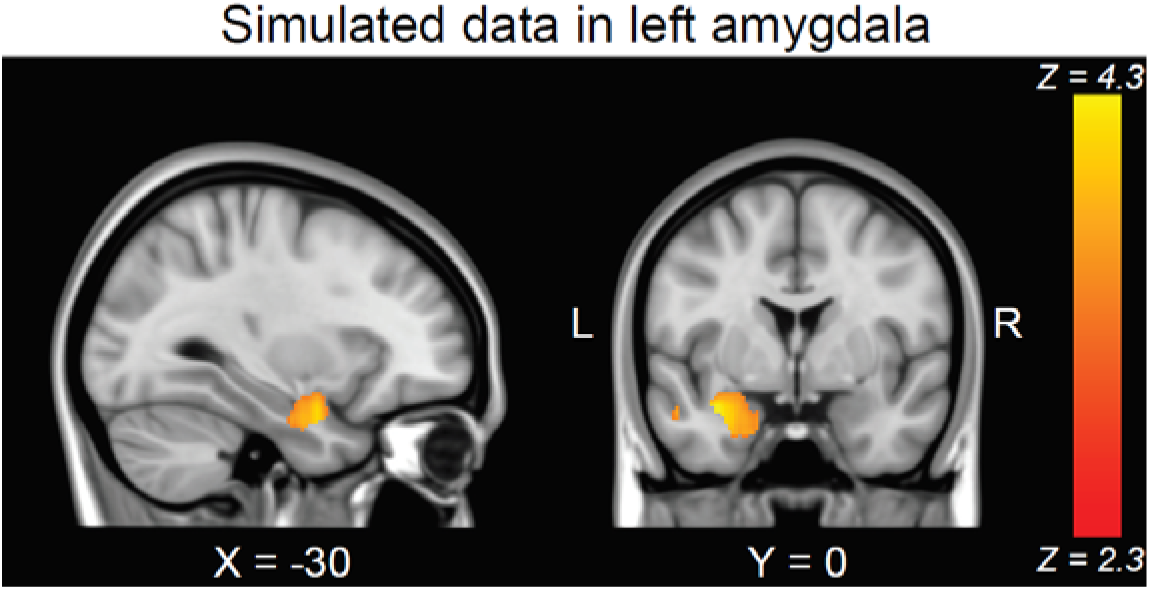
Data simulation Sub-cortical changes in the left amygdala were simulated in 10 participants. The SBM pipeline applied in CAT12 identified the simulated change effect. Data were projected from surfaces back to a voxel-based map for visualization in the current figure.

In an additional validation procedure, we reanalyzed our results with an additional analysis pipeline. Raw T1-weighted maps were preprocessed using FreeSurfer. Using this alternative analysis pipeline, we found similar results, including robust effects on the temporal cortical regions in addition to other cortical regions. It should be noted that the FreeSurfer pipeline can analyze only cortical surface, thus the analysis did not include subcortical regions of the amygdala and putamen.

The results of both validations indicated our analysis pipeline could reliably identify regional volumetric estimations. Using CAT12 provided an advantage by performing a longitudinal analysis of subcortical regions, including the amygdala which was pre-hypothesized and of great importance in the current work.

### Exploratory factor analyses: association with volumetric changes

As an exploratory analysis, we examined whether the volumetric brain changes were associated with the psychological constructs identified in our EFA, based on participants’ self-reports. In a previous version of this manuscript, we used principal component analysis (PCA) to identify our factors of interest, however this procedure is less appropriate than EFA, and is therefore not reported here. However, the results using PCA were fairly similar to the ones we found with EFA (previous version is available at:

https://www.biorxiv.org/content/10.1101/2020.09.08.285007v2).

We used two linear models to explain the variability in each of the factors, using the volumetric changes as our model features.

Overall, neither one of the factors was well associated with the volumetric changes (Factor 1 model: *R^2^* = 0.20, *F*(10,38) = 0.92, *p* = 0.522; Factor 2 model: *R^2^* = 0.22, *F*(10,38) = 1.10, *p* = 0.383). Examining the contribution of individual ROIs within the models (measured as the significance of the *β* estimates), did not reveal a significant association with the factors for any one of the ROIs (*p*_adj_. > 0.05; FDR correction by the number of features in the model). Adding the two factors as covariates to the linear models examining the volumetric change effect, also revealed no significant contribution of the factor to the models.

Thus, in our work we could not identify a clear association between the behavioral data and volumetric changes in our detected ROIs.

### Volumetric change correlation patterns

Finally, we examined correlation patterns of the volumetric change for all brain regions aiming to identify shared change patterns across multiple ROIs. Hierarchical clustering of the correlation matrices revealed different patterns between the two groups (Supplementary figure 4). In the test group, three principal groups of clusters could be identified - the first included the palladium, hippocampus, amygdala, putamen, insula (all bilaterally), and right anterior cingulate cortex, the second cluster included mostly occipito-temporal cortical and subcortical nuclei, and the third included highly correlated regions of the frontal, parietal and occipital cortices. All regions that passed the statistical threshold of the SBM analysis (Table 1), except for the left fusiform gyrus, were grouped closely together within the first two clusters, and showed low to negative correlations the ROIs of the third cluster. This analysis could suggest that the origin of the volumetric change observed in the regions of the two clusters might be different. The regions that appear in cluster 1 are often reported in the studies that explore brain changes following stress, anxiety or traumatic events^14–17^, while the regions of the second cluster are less associated with specific phenomena.

A different pattern was observed in the control group, where the correlation pattern demonstrated stronger volumetric synchrony (Supplementary figure 4b). This could suggest that changes in the control group were much more affected by within-participant effects, rather than an exogenous effect (which could be the outbreak and lockdown in the test-group). It is important to note in this context that in addition to the experience of the lockdown, the participants in the test group also had longer time gaps between the two scanning sessions, which might provide an alternative explanation for the stronger homogeneity in volumetric changes within the control group.

**Supplementary figure 4.**
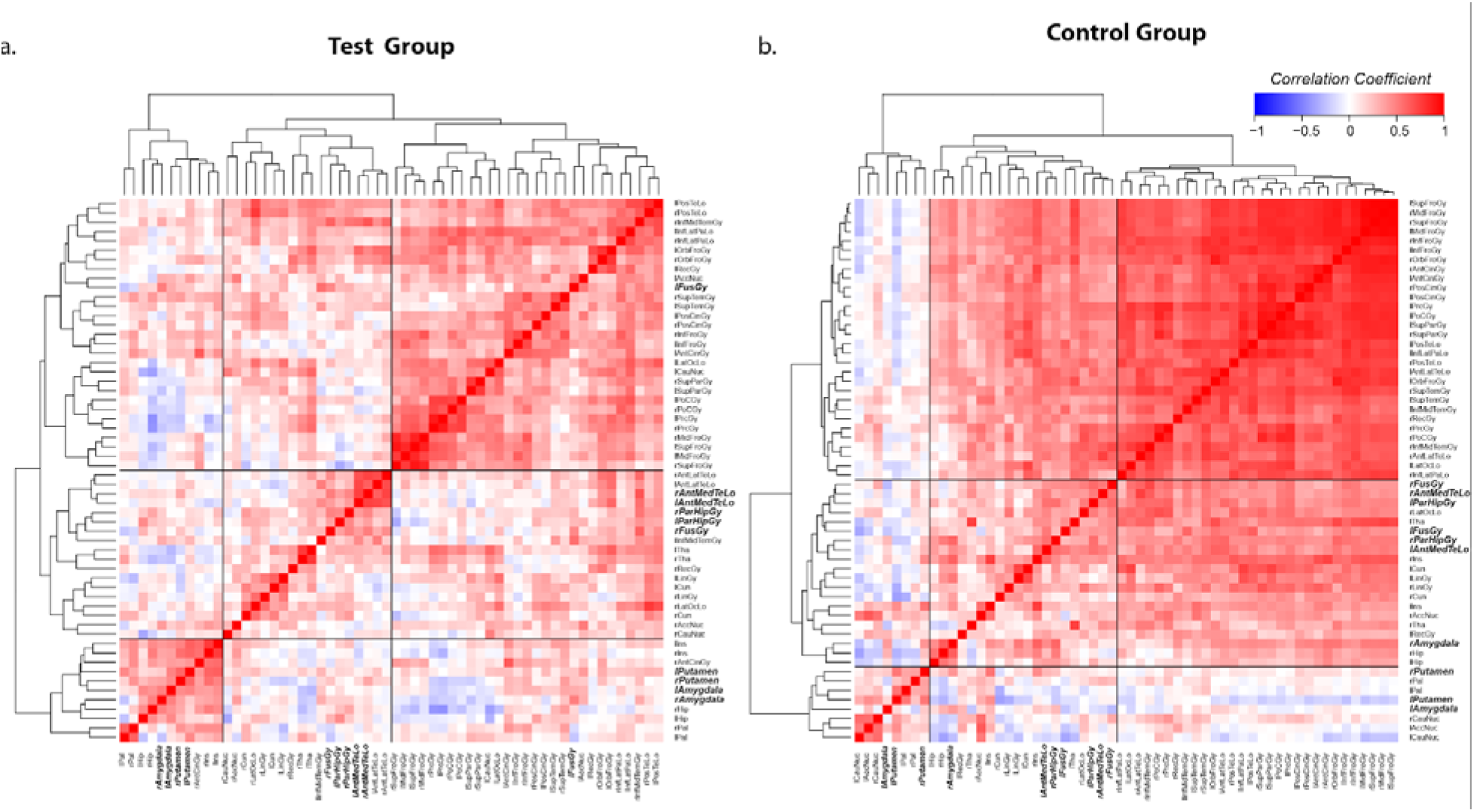
Volumetric changes correlation matrices Correlation coefficients were calculated between the volumetric change values for all ROI pairs, analyzed separately for the test **(a)** and the control group **(b)**, clustered according to Euclidean distances into dendrograms. In the test group, the first and second cluster contained the subcortical nuclei (amygdala and putamen) and temporal ROI which were found significant in the SBM interaction analysis, respectively (highlighted in italic bold font). In the control group, a more homogeneous change pattern was observed with more robust correlation coefficients between the ROIs. Pearson correlation coefficients are represented by the color scheme.

